# Differential Adaptation to Visual Motion Allows Robust Encoding of Optic Flow in the Dragonfly

**DOI:** 10.1101/496588

**Authors:** B. J. E. Evans, D. C. O’Carroll, J. M. Fabian, S. D. Wiederman

## Abstract

1

An important task for any aerial creature is the ability to ascertain their own movement (egomotion) through their environment. Neurons thought to underlie this behaviour have been well-characterised in many insect models including flies, moths and bees. However, dragonfly wide-field motion pathways remain undescribed. Some species of Dragonflies, such as *Hemicordulia tau,* engage in hawking behaviour, hovering in a single area for extended periods of time whilst also engaging in fast-moving patrols and highly dynamic pursuits of prey and conspecifics. These varied flight behaviours place very different constraints on establishing ego-motion from optic flow cues hinting at a sophisticated wide-field motion analysis system capable of detecting both fast and slow motion.

We characterised wide-field motion sensitive neurons via intracellular recordings in *Hemicordulia* dragonflies finding similar properties to those found in other species. We found that the spatial and temporal tuning properties of these neurons were broadly similar but differed significantly in their adaptation to sustained motion. We categorised a total of three different subclasses, finding differences between subclasses in their motion adaptation and response to the broadband statistics of natural images. The differences found correspond well with the dynamics of the varied behavioural tasks hawking dragonflies perform. These findings may underpin the exquisite flight behaviours found in dragonflies. They also hint at the need for the great complexity seen in dragonfly early visual processing.

**Significance Statement:** Understanding how animals navigate the world is an inherently difficult and interesting problem. Insect models have elucidated the neuronal mechanisms, which underpin this process. Neurons that encode wide-field motion have been studied previously in insects such as flies, hawkmoths and butterflies. Dragonflies exhibit complex aerobatic behaviours such as hovering, patrolling and aerial combat but little is known of their optic physiology. Moreover, dragonflies lack multimodal inputs (such as halteres in flies), which help enable diverse behaviours. The present study characterises wide-field motion sensitive neurons in the dragonfly. We find that wide-field motion sensitive neurons in dragonflies exhibit multiple subtypes, differentiated by their motion adaptation enabling encoding of a broad range of velocities independent of background contrast.

## 3 Introduction

Flying insects live in complex and varied 3-dimensional environments and display diverse flight behaviour, from near stationary hovering, to territorial patrolling and rapid pursuits of prey or conspecifics. This diversity places conflicting demands on the neuronal networks underlying self-motion detection. Neurons that respond robustly to patterns of wide-field motion have been extensively studied in several insect groups, including Dipteran flies (Hausen 1982, Hausen & Egelhaaf 1989), moths (Wicklein & Varju 1999, Theobald et al., 2010, Stöckl et al., 2016) and bees (DeVoe et al., 1982, Ibbotson 1991, Mertes et al., 2014). Typified by Lobula Plate Tangential Cells (LPTCs) of Dipteran flies, these neurons take input from local elementary motion detection (EMDs) elements located in the medulla (Borst et al 2010) and employ local correlation of spatially separated inputs with asymmetric delay mechanisms, consistent with influential computational motion models (Hassenstein & Reichardt 1956, Barlow & Levick 1965, Gruntman et al., 2018). Such neurons are tuned to specific spatial and temporal frequency ranges by their underlying spatial sampling and temporal delay filters. Because this places fundamental limitations on the velocity range of motion that neurons can individually encode, insects have evolved strategies for motion analysis that match their distinctive behaviour. For example, diurnal and nocturnal hawkmoths are precise hoverers when flower feeding and use wide field motion sensitive neurons specialized for such slow velocities (O’Carroll et al., 1996, 1997, Wicklein & Varju 1999, Theobald et al., 2010, Stöckl et al., 2016, 2017). By contrast, fast flying butterflies and bees show tuning to higher image speeds (Ibbotson 1991, O’Carroll et al., 1996).

In Dipteran flies, the conflicting demands of diverse flight modes that involve switches between slow hovering to high speed pursuit flight are in part met by multimodal integration of fast input to descending visual pathways from the ocelli (Parsons et al., 2006) and specialized hindwing mechanosensory organs (halteres) that detect rapid accelerations, allowing compound eye neurons (the LPTCs) to focus on slower motion (Hengstenberg 1991).

Although dragonflies have recently emerged as an important model for studying visual target tracking, very little is known about their neural tuning to wide-field motion. Dragonflies exhibit a similar behavioural repertoire to Dipterans, but have a lower wingbeat frequency, and lack specialized halteres for detecting gyroscopic forces. As an essentially visual creature, how do dragonflies encode the large velocity ranges demanded by their behavior? One potential strategy is to process the same retinal input using parallel pathways employing spatiotemporal filters tuned to different speed ranges, as seen in mammals (Movshon & Newsome 1996, Nassi & Callaway 2008). In many insects, however, replicating such parallel pathways may be constrained by their size and weight. Indeed, in species studied to date, motion tuning at a behavioral level appears to reflect a single common EMD mechanism (Buchner 1976). Nevertheless, we hypothesize that parallel processing may be viable for dragonflies, with among the largest eyes and brain of extant insects. Alternatively, useful coding of different speed ranges may result from additional downstream processing. Motion adaptation, for example, can improve velocity contrast via relief from saturation (Maddess & Laughlin 1985; Barnett et al., 2010) and on a timescale similar to the stimulus response (Nordström et al., 2011). It also improves velocity encoding of natural images (Shoemaker et al., 2005, Straw et al., 2008, Barnett et al., 2010) and enhances differentiation between foreground and background features (Li et al., 2017).

We tested these two alternative strategies by recording from widefield motion-sensitive neurons in the lobula of dragonflies. We found evidence for several unique subclasses of widefield motion-sensitive neurons, some of which differ substantially from their counterparts in other species. We found evidence that these neurons likely share common input pathways (i.e. using the same EMD inputs) but differ radically in their adaptation to image motion. This differential motion-adaptation tunes otherwise similar neurons to significantly different velocity ranges, providing very robust encoding of motion over several decades of image speed.

## 4 Materials and Methods

### 4.1 Electrophysiology

93 wild-caught, dragonflies (*Hemicordulid*) were immobilized with a 1:1 beeswax and rosin mixture, with the head tilted forward to access the posterior surface. A hole was cut above the brain to gain access to the lobula and lateral midbrain, but the preparation was otherwise left with the perineural sheath and overlying haemolymph sacs intact. We penetrated the sheath and recorded intracellularly using strong aluminosilicate micropipettes, pulled on a Sutter Instruments P-97 puller and backfilled either with KCl (2M, electrode tip resistance typically 50-150 MΩ) or 4% Lucifer Yellow solution in 0.1M LiCl.

### 4.2 Visual Stimuli

We presented stimuli on high definition LCD monitors (120 -165 Hz). The animal was placed 20 cm away and centred on the visual midline. Contrast stimuli were presented at screen centre to minimize off-axis artefacts. The display was projection distorted using OpenGL to ensure each 1° onscreen was 1° from the animal’s perspective. The visual field was 104° by 58.5°. All temporal frequencies tested were limited to one quarter of the monitors frame rate. Stimulus scripts were written using MATLAB’s Psychtoolbox and integrated into the data acquisition system.

To classify neurons as widefield motion sensitive, a sequence of characterising stimuli were presented to the dragonfly. These included a gyrated, randomly generated texel pattern (1°), grey to black and grey to white full screen flicker (White – 338 cd/m^2^, Black 0.5 cd/m^2^), moving edges (up, down, left and right, 25°/s), moving bars (2° width, up, down, left and right, 25°/s) and a square-wave grating pattern moving up, down left and right (0.025 cycles/°, 6.25Hz). Neurons were categorised as widefield motion sensitive based on robust responses to the gyrated texel pattern and square-wave gratings. Following, sinusoidal gratings were presented to dragonflies which had a linear increase in contrast for 1 s (0 to 0.25, Weber) followed by a 1 s exponential rise (0.25 to 1).

### 4.3 Neuroanatomy

The morphology of a widefield motion-sensitive neuron was visualized by intracellular labelling with Lucifer Yellow (Figure 1a, b, c). Iontophoresis was achieved by passing 1nA negative current through electrodes tip-filled with Lucifer Yellow for 12 minutes. Brains were then carefully dissected, fixed overnight in 4% paraformaldehyde at 4°C, dehydrated in ethanol series (70%, 90%, 100%, 100%), cleared in methyl salicylate and mounted using Permount on a slide using three spacer rings and covered with a cover slip for imaging. The sample was scanned using a confocal microscope using a 10x objective and the 3D slices reconstructed using Neutube.

**Figure 1:**
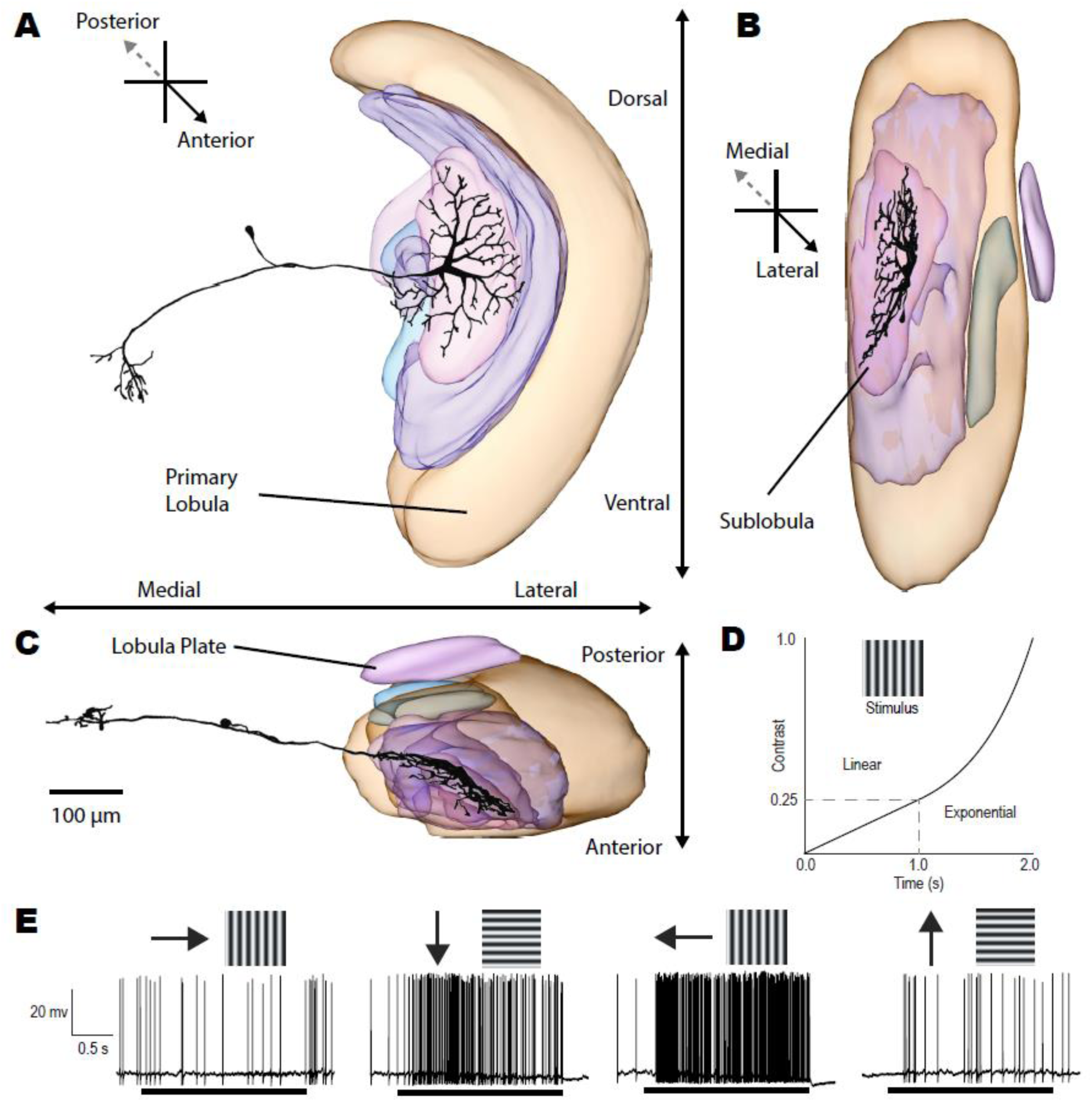
**A,** Anterior, **B,** Longitudinal and **C,** Horizontal projections for a 3D model reconstruction of a dragonfly lobula tangential cell (LTC). **D,** description of the waveform of the contrast ramp used to characterise response tuning and contrast sensitivity in LTC neurons. At cessation of a pre-stimulus period, the contrast rises linearly for a 1 second, from 0 to a contrast of 0.25. Then for the remaining 1 second, the contrast continues to rise exponentially from 0.25 to a final contrast of 1.0. **E,** Example responses to sinusoidal gratings in four directions for a direction opponent LTC. Black bars indicate the period of the 2 second contrast ramp. Direction opponent cells are inhibited by motion in the ‘anti-preferred’ direction (left) while responding strongly when stimulated by the opposite direction of motion.

### 4.4 Experimental Design and Statistical Analysis

All analysis was completed in MATLAB. Spike-counting was done using a custom-written spike-counting script. Curve fits used MATLABs in-built curve-fitting tools. To find peaks of tuning curves, repeated measures were averaged followed by the application of a 5-point moving average filter to smooth data before finding the maximum. All statistical tests were either paired-sample non-parametric tests (Wilcoxon sign test, for paired data) or two-sample non-parametric tests (Mann-Whitney U test, for unpaired data) with appropriate multiple-comparison corrections (Bonferroni). All means are calculated from biological replicates (i.e. repeated measurements from identified neurons in different animals). Each biological replicate represents the mean of between 1 and 5 technical replicates. P values are reported as raw numbers in text if significant (unmarked otherwise) or as < 0.0001 if sufficiently small. Box and whisker plots represent the 75^th^, 50^th^ and 25^th^ quartiles (lines) with raw data shown.

## 5 Results

### 5.1 Neuroanatomical characterisation

While we were not able to anatomically identify every neuron that we recorded from in the midbrain/lobula complex in the dragonfly, we nevertheless stained a subset of these neurons, typified by the example in Figure 1. This confirms a similar general organisation to that in Diptera and several other insect orders (Hausen 1982, Egelhaaf 1989, Fabian 2017), where optic flow is integrated within specialised subregions of the 3^rd^ optic ganglion (the lobula complex) by a set of *tangential* neurons, the well-studied wide-field motion sensitive ‘Lobula Plate Tangential Cells’ (Hausen 1982). These neurons have input dendrites that integrate tangentially across arrays of retinotopically-organised inputs from underlying local motion detectors (presumptive EMDs) at earlier stages of visual processing. Figure 1A-C shows the reconstructed morphology of a dragonfly neuron that exhibited wide-field motion sensitivity. The overall morphology of these neurons strongly resembles their Dipteran counterparts, with tree-like input arborisations within the lobula complex, and outputs in the lateral midbrain. However, as with this individual example, several neurons described in this study have their inputs originating solely from a deep neuropil on the anterior side of the lobula, similar to the ‘sublobula’ identified in bees (Devoe et al., 1982, Strausfeld 2005, Strausfeld et al., 2006), rather than from a posterior Lobula Plate. Until the homologies between these different lobula subregions with their counterparts in other insect groups are more clearly identified, we label these neurons more generally as ‘Lobula Tangential Cells’ (LTCs).

### 5.2 Direction selectivity and opponency

We tested the motion sensitivity of LTCs using sinusoidal gratings drifted in eight directions (presented randomly in 45° increments). Each grating was displayed as a ‘contrast ramp’ with a nonlinear increase in contrast from zero over a 2 s period (Figure 1D). This stimulus avoids onset-transients inherent with step changes in contrast. The ramp also weights more time around the threshold contrast values whilst still providing a stimulus that contains the entire contrast range (O’Carroll et al., 1996, 1997). Figure 1e shows an individual neuron’s spiking response to four directions of motion. This LTC response exhibits clear direction opponency, with excitation to preferred motion and inhibition to the anti-preferred direction of motion. The response time course in these neurons typically shows high initial sensitivity to low contrast, indicated by a rapid rise in firing rate. The ramp responses show variable degrees of saturation as the contrast continues to increase and in many cases a subsequent decrease in response due to temporal adaptation.

We measured LTC directionality by presenting a gyrated random (binary) texture pattern (Figure 2A). This continuous stimulus is composed of a broad range of spatial and temporal frequencies and permits precise calculation of direction selectivity. The full-screen texture was gyrated (not rotated) in a clockwise and counter-clockwise direction for two complete cycles. In direction opponent neurons, this elicits a sinusoidal pattern of excitation and inhibition (Figure 2B). We thus fitted a sinusoid (Figure 2C) to the instantaneous spike rate (inverse inter-spike interval). Responses to clockwise and counter-clockwise rotations were then averaged to eliminate any phase-lag due to response latency. The maximum and minimum values of the fitted sinusoid were taken as the neuron’s preferred (R_p_) and anti-preferred (R_a_) responses. We then defined a direction opponency metric relative to the spontaneous activity level (R_s_) as:

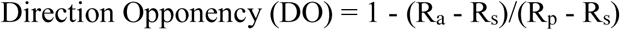

**Figure 2:**
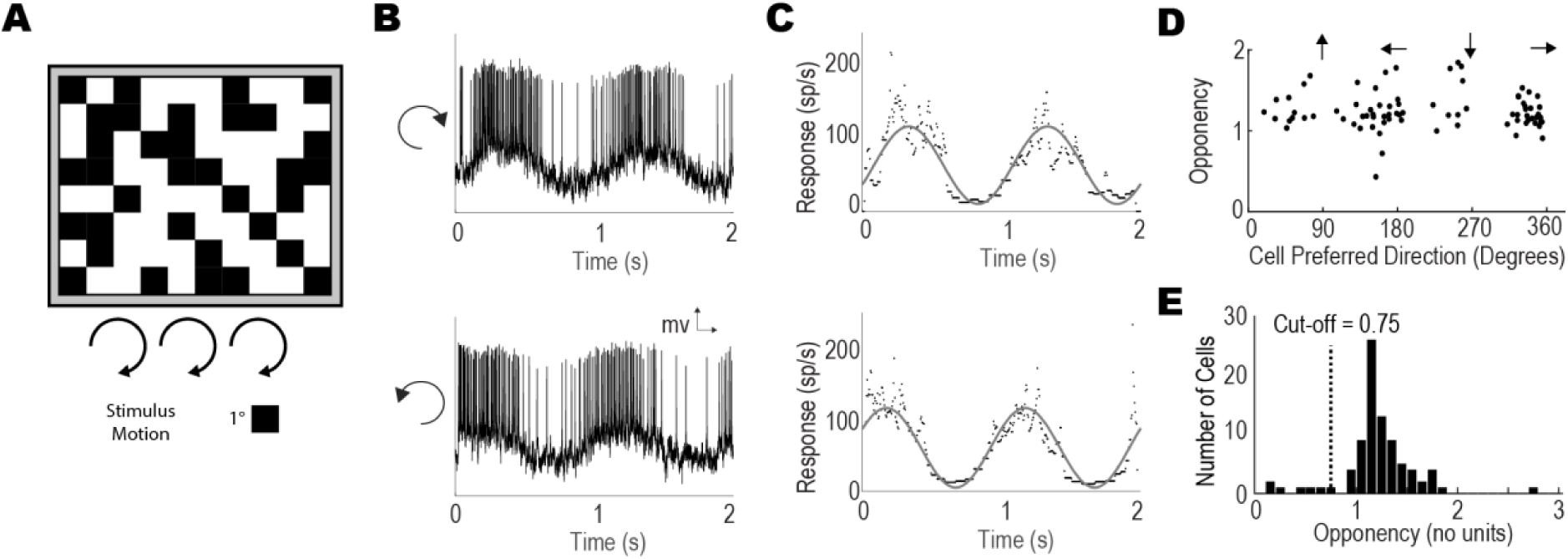
**A** Moving texel patterns (1° texels) at a constant orientation were presented to the dragonfly whilst recording from widefield motion sensitive neurons. The pattern was gyrated twice in a clockwise circle (not rotated) and then twice anti-clockwise at 50°/s. **B.** An example spike trace during texel pattern stimulus shows a clear periodic response corresponding to the texel pattern direction of motion. **C.** The inverse interspike interval (ISI) reveals spiking activity to the stimulus, which is fitted with a sinusoidal curve for both a clockwise (top) and counter-clockwise (bottom) texel gyration. This curve fit provides both the peak response phase (i.e. preferred direction) and response characteristic at the preferred and anti-preferred direction establishing the neuron’s directionality **D.** Each point indicates the neuron’s preferred direction and DO. Dragonfly LTCs exhibit preference for four directions (progressive, regressive, upward, downward). **E.** A histogram showing the direction selectivity (in total neurons) of the widefield-motion sensitive neurons.

We used this basic stimulus to characterise the preferred direction direction selectivity in 93 neurons (recorded from the dragonfly lobula complex) that gave strong responses to wide-field motion (Figure 2D). These opponent LTCs exhibit both vertical and horizontal preferred directions, clustering around all four cardinal directions (i.e. left, right, up and down). A similar alignment of neuronal sensitivity to the different directional components of ego-motion are also observed in the frontal visual fields of LPTCs in other insect species (Strausfeld & Lee 1991, Krapp & Hengstenberg 1996).

This initial neuron selection was likely biased by repeated recordings from this stereotyped location in the optic lobes where we had previously located direction selective cells. Hence the resulting distribution of DO values probably underestimates the number of neurons that are not direction selective. Nevertheless, while the recorded neurons displayed a large range of DO (from near zero to 2.8), a histogram of DO reveals a clear peak in the distribution above 1.0, indicating that most of the neurons were strongly direction opponent (Figure 2E). Strong direction selectivity and opponency is a characteristic of many of the LPTC neurons seen in other taxa, so the role of the more weakly directional neurons from this group in optical flow analysis is unclear at this stage. We therefore limit our subsequent analysis in this paper to a subset of recorded neurons with strong direction selectivity (DO>0.75), which are more likely to be analogues or homologues of the LPTC neurons in other species (Figure 2E).

### 5.3 Spatial and Temporal Tuning

In many animal models, motion sensitive neurons exhibit distinctive tuning to both the spatial and temporal frequency of drifted gratings (Hausen 1982, Devoe et al., 1982, Arenz et al., 2017). Here we established LTC spatiotemporal tuning by presenting two series of sinusoidal gratings. To establish spatial tuning, we presented a randomised series of 30 gratings of varying spatial frequencies logarithmically spaced between 0.01 and 1 cycles/°, using a fixed temporal frequency (5 Hz). To establish temporal tuning we presented a series with varying temporal frequencies logarithmically spaced between 0.1 and 30 Hz, using a fixed spatial frequency (0.1 cpd). In each case the grating stimulus was a ramp of contrast as previously described (Figure 1D).

Figure 3 shows data for a single neuron that exemplifies a subset of LTCs that show consistency in their spatial and temporal tuning across time, and thus to increasing contrast as the ramp stimulus progresses. This particular neuron gave mixed mode responses, with spikes that ride on graded depolarization when the stimulus was excitatory (Figure 3A). Separate quantitative analysis of such mixed-mode responses revealed general consistency between the graded and spiking responses across a wide range of stimulus conditions. Since other neurons showed only biphasic (axonal) action potentials (e.g. Figure 1) we thus limit our subsequent analysis to the spiking component of the activity. At very low spatial frequencies, responses are often phase-locked to the 1^st^ harmonic of the stimulus waveform (i.e. the original 5 Hz frequency), particularly at high contrast (e.g. Figure 3A top trace). This phase-locking is not evident, however at higher spatial frequencies. Stimulus conditions that elicit the strongest responses during presentation of the grating also lead to a strong rebound response on motion cessation, i.e. a motion after effect (Anstisa et al 1998, Nordstrom et al., 2009) very evident in the graded response, but also in a reduction in spike firing rate in the post-stimulus period.

**Figure 3:**
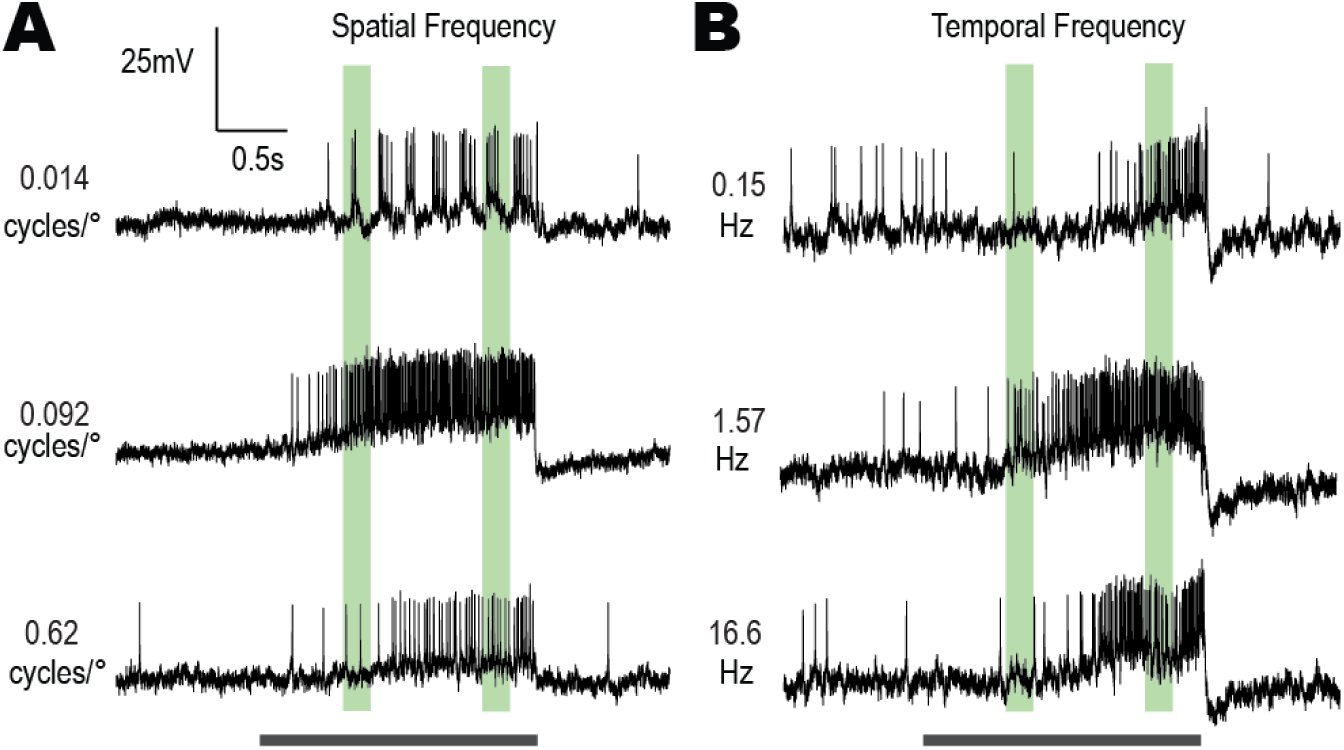
Responses from a neuron that exemplifies a subset of LTCs that show strong consistency in their spatial and temporal tuning over time (and thus increasing contrast). The period of the contrast ramp stimulus is indicated by the black bar. Green bars represent early and late window periods used for later analysis. **A,** raw responses to 3 different spatial frequencies, 0.014, 0.092 and 0.62 cycles/° using gratings with a temporal frequency of 5 Hz. **B,** corresponding raw responses to 3 different temporal frequencies (0.15, 1.57 and 16.6 Hz) at a constant spatial frequency of 0.1 cycles/°.

### 5.4 Temporal Adaptation and Neuron Classification

Do the direction opponent LTCs show evidence of more than one class? 3-dimensional plots of different LTC responses over time show remarkably different characteristics, particularly in the temporal frequency domain (Figure 4A-C, note colour is the change from spontaneous activity). Such plots reveal responses over a broader range of frequencies as time progresses, reflecting recruitment of activity for less-optimal stimuli as grating contrast increases. This initial broadening was observed in all neurons recorded, although we found that they varied widely both in their initial contrast thresholds and in their subsequent responses due to differences in temporal adaptation. To account for these differences and yet still compare initial (weakly adapted) with later motion-adapted responses we also derived tuning curves (Figure 4D-F) based on two 200 ms duration analysis windows (as indicated in Figure 3). The *early* window (black line) starts when responses exceed the spontaneous activity by 2 standard deviations to an optimal stimulus. A *late* window (red line) begins 250 ms prior to the end of the ramp stimulus.

**Figure 4:**
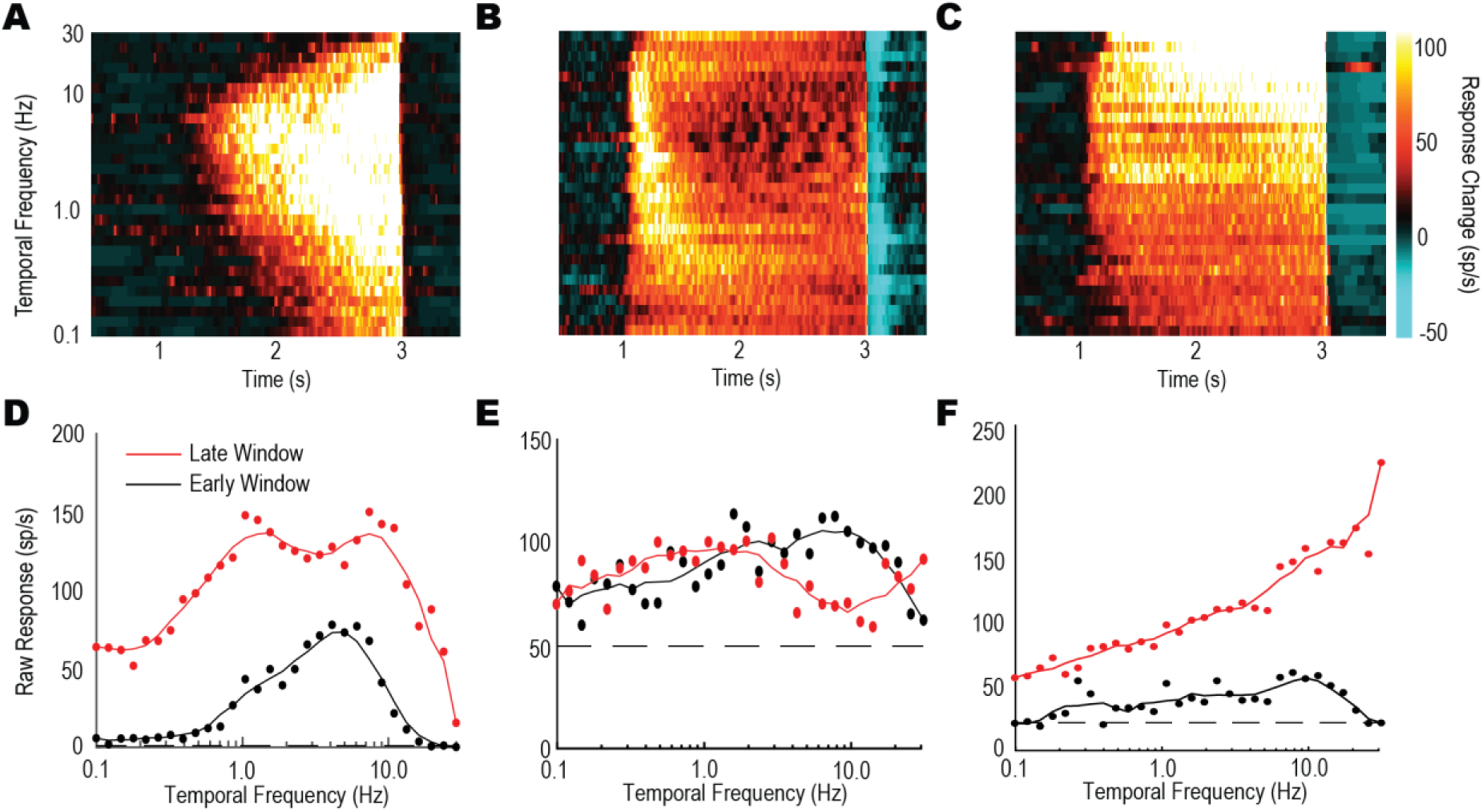
**A-C,** 3-dimensional plots of the spike rate above spontaneous activity over time in response to contrast ramps at 30 different temporal frequencies (at a spatial frequency of 0.1 cycles/°) for examples of 3 different putative neuron sub-classes; (**A** slow adapting tangential cells (SATC), **B** selective fast adapting tangential cells (SFATC), **C** fast adapting tangential cells (FATC). Each plot shows the spiking activity during the 2 second contrast-ramp gratings (colour coded for the inverse interspike interval i.e. spike/s). **D-F** Temporal tuning curves derived by averaging responses from plots as in **A**-**C** within short (200ms) windows either early in the ramp (low contrast, black line) or late (high contrast, red line).

In the first example LTC (Figure 4A, D), responses continue to increase with higher contrast, saturating across a range of temporal frequencies for the near-optimal spatial frequency used here (0.1 cpd), but with no obvious shift in the location of the optimum. Consistency of spatial and temporal optima, and robustness of the basic shape with respect to adaptation state of neurons (as in Figure 4A, D) is also observed in *Dipteran* LPTCs (Hausen 1982, Harris et al., 1999). However, not all LTCs exhibited such LPTC-like tuning properties, however. Figure 4B and Figure 4C show data for 2 additional neurons, highlighting large variation in the evolution of responses over time for different temporal frequencies (though all still exhibiting direction selective responses to wide-field motion). We first defined a subclass of neurons like that in Figure 4A and Figure 4D, showing little change in their tuning properties over time. We termed these *slow adapting tangential cells* (SATCs). Qualitatively, the other neurons (Figure 4B, C) exhibited much stronger motion-adaptation, though in different ways. A second subclass exhibit very strong motion adaptation at their initial preferred temporal frequency, so that even high contrasts at the end of the ramp at such frequencies elicit only weak responses (Figure 4B, E). This gives rise to a distinctive ‘notch’ in the centre of the late window temporal tuning (Figure 4E) which is also evident from the dark region in the 3 dimensional plot (Figure 4B). Despite this adaptation at the initial optimum frequency, these neurons retain strong responses at both higher and lower temporal frequencies. We refer to these as *selective fast adapting tangential cells* (SFATCs) to account for the selective nature of the motion adaptation by intermediate frequencies. A third group of neurons (Figure 4C, F) strongly adapted to low temporal frequency stimuli, shifting their most robust responses to higher temporal frequencies over time, reaching the limits of frequencies possible with our display. We termed this subclass *fast adapting tangential cells* (FATCs).

### 5.5 Classification of Subclasses of LTC

Whilst the differences between individual neurons as illustrated in Figure 4 is observed in the examples selected, formalizing this subdivision across our population of recorded neurons based on quantitative measures is more difficult. When we plotted 3-dimensional plots of the spike rate over time in response to contrast ramps at 30 different temporal and spatial frequencies for a larger group of neurons, the patterns described above for the different subclasses was obvious in many, but not all cells (Figure 5). Each panel in this figure uses the same colour lookup table to indicate spike rate, so overall responsiveness can be compared across neurons. In a subset of neurons in which we were able to record mixed mode responses, we also examined the graded response component (data not shown). We found these to be qualitatively similar to the spiking data, so our subsequent analysis is limited to spiking responses. For each individual cell we have provided panels for the spatial and temporal tuning over time (paired vertically).

**Figure 5:**
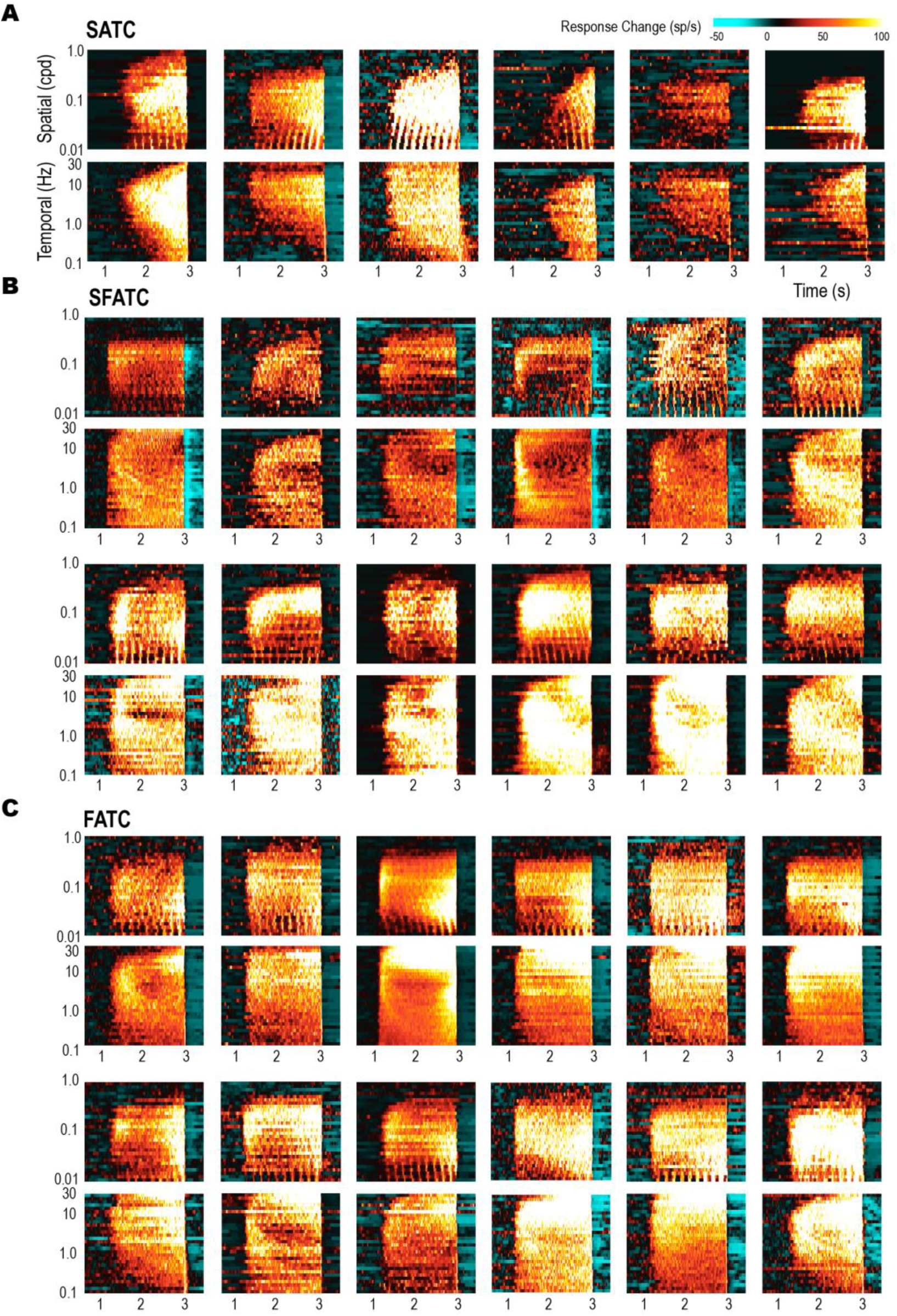
3-dimensional plots of individual LTCs forming putative subclasses. The change in spike rate (spontaneous is subtracted) over time is plotted in response to contrast ramps at 30 different spatial and temporal frequencies. Plots are arranged in pairs for each individual neuron, with responses to varying spatial frequencies (temporal frequency of 5Hz) arranged above that for temporal frequencies (spatial frequency of 0.1 cycles/°). Each plot shows the spike rate during the 2 s stimulus duration (contrast-ramp gratings) which includes a 1 s pre-stimulus and 1 s post stimulus period. **A**, Slow adapting tangential cells (SATCs, 6 examples shown) exhibit LPTC-like tuning, that broadens over time (increasing contrast) with little shift in the optimal tuning **B**, Selective fast adapting tangential cells (SFATCs, 12 examples shown) with their readily identifiable adapted ‘notch’ forming at the earlier optimal tuning. **C,** Fast adapting tangential cells (FATCs, 12 examples shown) revealing strong shifts in spatial and temporal frequency tuning over time.

The initial division into subclasses is supported by consistent and characteristic differences in the pattern of motion adaptation, as described for the examples provided in Figure 4. Most SATCs (Figure 5A) show little motion adaptation, responding more strongly at any given frequency as pattern contrast increases and thus to a broader range of frequencies as time progresses. SFATCs and FATCs (Figure 5B, C) both show more complex time courses, with darker coloured areas often following the recruitment of responses at the initial optima in different bands of both spatial and temporal frequency.

The SFATC and FATC subclasses have more complex interactions between motion adaptation (leading to weaker responses) and increasing contrast (leading to stronger responses) during the course of the contrast ramp. This diverse group may ultimately prove to represent a spectrum of response characteristics. Nevertheless, to permit comparison of other response parameters between them, we sorted all recorded neurons into the 3 groups as follows. First, after visual inspection of the late versus early windowed data, neurons with clearly 2-peaked adapted temporal tuning (i.e. the ‘adaptation notch’ identified in Figure 4E) were considered to be SFATCs. For the remaining 42 neurons, which all exhibited a single peak in temporal frequency tuning in both the early and late windows, we quantified the shift in optimum as a fold change from the original value described in the log-domain (thus 0 represents no change and 1 a ten-fold change). The distribution of these temporal frequency ‘peak change’ values is shown in Figure 6A. From Figure 6A it is difficult to see a clear cut-off point between the two cell subtypes and it is likely that the two populations overlap using this metric. Despite this, by choosing an arbitrary threshold (0.32 log units, ∼2.07-fold change), the two subtypes were well separated when compared qualitatively (SATC green bars, FATC red bars). Such shifts in temporal frequency optima over time (and therefore increasing contrast) is a novel observation in insect, wide-field motion sensitive neurons, with responses observed in Dipteran LPTCs more like those classified here as SATCs (Figure 6A, green bars, < 0.32 log peak change).

**Figure 6:**
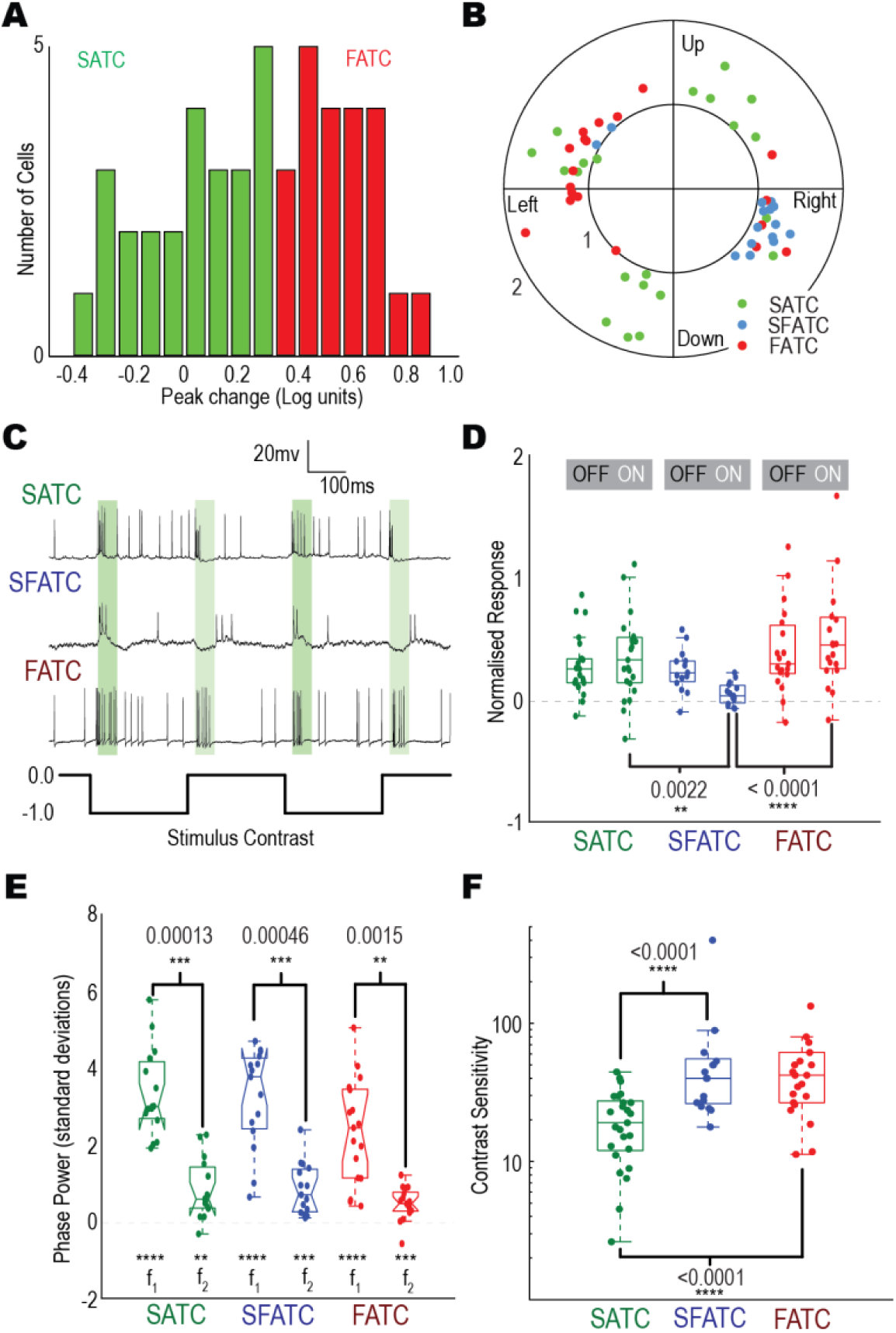
**A,** Histogram depicting the logarithmic shift in temporal frequency optimum in the group of LTC neurons identified as putative SATCs and FATCs. **B,** Polar plot of direction tuning (angle; across up down, left and right) values plotted against direction opponency (magnitude, where DO>0.75, see Figure 2). FATC and SFATC cells appear to code horizontal motion directions only, while SATCs include individual neurons with peak responses towards any of the 4 cardinal directions. **C,** Raw traces from each subclass of LTC when stimulated by full-screen flicker (grey to black, 0.0 to -0.99 Weber contrast). Green bars indicate window periods (20 to 70 ms) for analysis. These individual SATC and FATC cells exhibit full-wave rectification (increased responses to both ON and OFF increments) while SFATCs were inhibited to ON flicker. **D,** Boxplot distributions of windowed responses to full screen flicker (0.73 and -0.99 Weber contrast), following either luminance increments (ON) or decrements (OFF). SFATC cells show no response to ON flicker, while FATC and SATCs generally show responses to both ON and OFF components. **E,** boxplot distributions of the fundamental frequency (f_1_) and second harmonic (f_2_) responses estimated in the late part of ramp responses at low spatial frequencies (mean of results from 0.137-0.260 cycles/°), at a temporal frequency of 5 Hz. All 3 classes of neurons show significant modulation at both f_1_ and f_2_ **F,** Boxplot distributions of contrast sensitivity (to a drifting grating) based on the contrast required to evoke a neuronal response (2x the standard deviation of the spontaneous activity). SFATC & FATC cells exhibit significantly higher peak contrast sensitivity than SATCs (Mann-Whitney U test, n=65).

### 5.6 Differences in response sensitivity and tuning across subclasses of LTC

Does our classification of these cells into several subclasses (based primarily on differences in their temporal adaptation properties) correlate with other physiological response attributes? Figure 6B shows the direction tuning data as defined earlier (Figure 2) replotted for the different subclasses of LTCs on polar axes. Neurons selective for horizontal motion include examples from all 3 subclasses, but interestingly all of the LTCs with vertical preferred directions (i.e. sensitive for upwards or downwards motion) fell into the SATC subclass. Given their frontal receptive fields and response characteristics resemble those of Dipteran LPTCs, these horizontal and vertical sensitive SATC neurons may indeed be the dragonfly equivalent of the VS (vertical system) and HS (horizontal system) neurons that play an important role in analysis of pitch, roll and yaw stimuli (Hausen & Egelhaaf, Krapp & Hengstenberg 1996).

In our initial characterisation, we noted that all LTCs give transient responses to full-screen (square wave) flicker at low temporal frequencies (Figure 6C). Much recent work on Dipteran LPTCs supports a model that integrates inputs to local motion detectors from separate ON and OFF pathways that originate in early visual processing (Borst & Helmstaedter 2015). By contrast, our own prior work suggests that motion detection in the feature-selective pathways of this same dragonfly species can be strongly selective for the OFF pathway (Wiederman & O’Carroll 2013). Do we see any clear segregation of these flicker components in the responses of different LTC subclasses? Figure 27C shows example data traces of individual LTCs in response to a 2 Hz full-screen flicker pattern. We analysed response windows 20 to 70 ms after the onset of each ON and OFF phase (shaded green regions) for the various LTC subclasses. Figure 6D shows these ON and OFF response components for our population of LTCs, separated into the subclasses. For both SATC (green) and FATC (red) responses are full-wave rectified, with similar spiking responses to both ON and OFF transitions. However, SFATCs responded predominantly to the OFF transition in the full-screen flicker stimulus with hyperpolarisation during ON stimuli (luminance intensity increases). This resulted in a statistically significant difference between the OFF and ON responses of SFATCs (n=15) and a statistically significant difference between SFATC ON responses and both SATC and FATC ON responses (Kruskal-Wallis with multiple comparisons, SATC n=23, SFATC n=15, FATC n=19). This result is somewhat at odds with our observation that this subclass, like the SATCs and FATCs, exhibit a clear phase-locked modulation of the response to very low spatial frequency grating pattern drifted in the preferred direction (as in Figure 3A). That is, in response to low spatial frequency motion, responses modulate at the grating’s temporal frequency (1^st^ harmonic, f_1_). However, the transient changes of full-screen flicker results in frequency-doubled, ‘breakthrough’ responses to both ON and OFF intensity changes (2^nd^ harmonic, f_2_).

To interrogate these phase-locked responses, we performed a fast Fourier transform on the mean response of five of the lowest spatial frequencies tested (0.0137-0.0259 cycles/°). This frequency of these low frequency motion responses (Figure 6E, SATC n=15, SFATC n=15, FATC n=18) shows that they are dominated by power at the fundamental temporal frequency of the grating (i.e. the first harmonic) with weaker but still statistically significant power (Kruskal-Wallis with multiple comparisons) at double the fundamental temporal frequency (i.e. the second harmonic), suggesting a fully rectified input (i.e. both ON and OFF responses) in all 3 subclasses of LTC.

A clear distinction between the SATC and FATC/SFATC classes is also evident from their contrast sensitivity. This is observed from the 3D plots in Figure 5, where many FATC and SFATC cells show an abrupt transition from the pre-stimulus (spontaneous) response level to strong excitation within a few hundred milliseconds of ramp onset. This indicates a very high sensitivity to very low contrast across a large range of frequencies. To quantify this sensitivity further (Figure 6F), we estimated the standard deviation of the spontaneous activity of the neuron when viewing a blank screen in the pre-stimulus period and then determined the time during the ramp at which the response exceeds 2x this value and back-calculated the contrast based on the ramp waveform. We took the inverse of this threshold value as the sensitivity. This method equivalent to the detectability criterion used in previous studies of contrast sensitivity in insect motion sensitive neurons (Dvorak et al 1980). Based on this criterion, all 3 neuron classes showed high contrast sensitivity, with peak values in the range from 20 (for SATCs) to 40 (for FATCs), similar to peak sensitivity values reported for other insect species (O’Carroll & Wiederman 2014). Nevertheless, peak contrast sensitivity was significantly higher in FATCs and SFATCs (SATC n=25, SFATC n=17, FATC n=22, Kruskal-Wallis with multiple comparisons test) that display clear motion adaptation during the ramp, suggesting that adaptation may help compensate for higher initial contrast gain to avoid saturation.

### 5.7 Spatial tuning

Figure 8 shows spatial tuning (temporal frequency = 5Hz) data averaged across all cells allocated to the SATC, SFATC and FATC subclasses. All three classes exhibit qualitatively similar spatial tuning in the early analysis window, before adaptation at higher contrasts begins to affect the curve shapes. The optimum is centred near 0.1 cycles/°, with high sensitivity extending responses to the lowest frequencies tested (0.01 cycles/°), but rolling off above 0.5 cycles/°. This roll-off is consistent with the theoretical predictions based on the interommatidial angle, which is a little below 1° for this species (Horridge 1978, Buchner 1976). In the late windows, however, higher contrast and differential adaptation as ramps progress alters the shape of spatial tuning curves among different cells in ways that reinforce our earlier segregation of the different LTC subclasses based on other parameters. Firstly, consistent with their contrast sensitivity (Figure 7E) SFATCs show stronger responses than SATCs in the early window and less response increase in the later window (i.e. as responses saturate). Apart from this saturation, the late and early window responses have similar shape in SFATCs, with little evidence of a ‘notch’ near the initial optimum as seen in the temporal frequency domain. Although the apparent centre of the distributions is similar for the averaged tuning data (Figure 7A) individual cells do show a weak increase in the optimum spatial frequency for the SFATC cells (Figure 7C). Interestingly, although our distinction between SATCs and FATCs was made solely on the basis of the shift in their temporal optima, the group we identified as FATCs clearly also show a difference in the shape of their spatial tuning as contrast increases, with a significant reduction in the optimum towards a 3-fold lower spatial frequency (Figure 7C, n=18, Kruskal-Wallis with multiple comparisons test).

**Figure 7:**
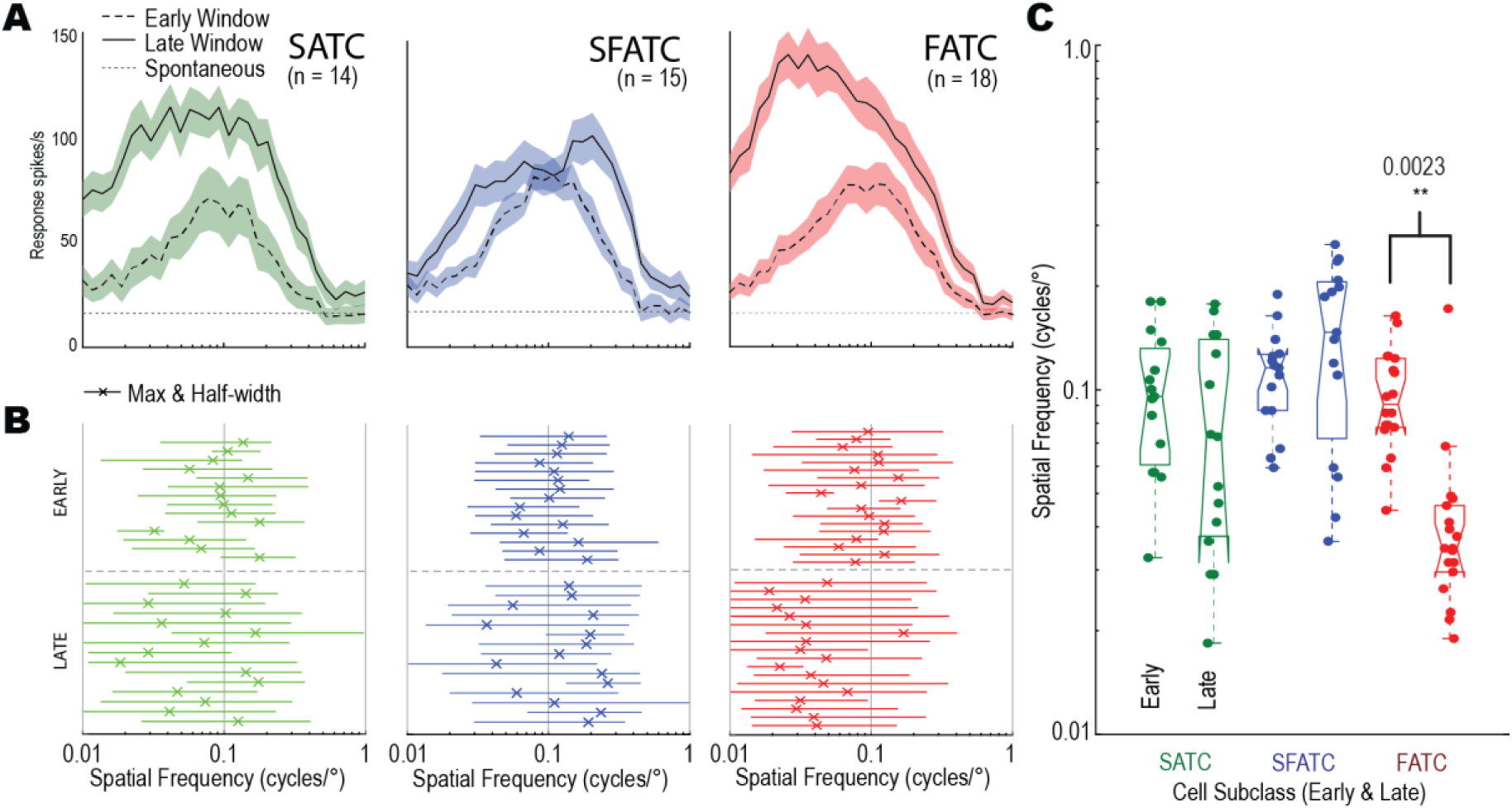
Spatial tuning of different LTC subclasses. **A**, Average spatial tuning for different cell classes in two response windows (early, late) as defined in Figure 4. Each plot shows the mean response as a line with the shaded area denoting standard error. The mean spontaneous across cells is indicated by the grey dotted line. FATC cells exhibit a clear shift towards lower spatial frequencies in their late window compared with SATCs & SFATCs. **B**, Individual neuron summary data showing the position of the peak in early/late windows (denoted by an X) with the full width at half maximum (width of responses greater than 50% of maximum response) indicated by horizontal lines. **C**, Changes in spatial frequency peaks shown as box plots. FATCs show significant decrease in peak spatial tuning.

**Figure 8:**
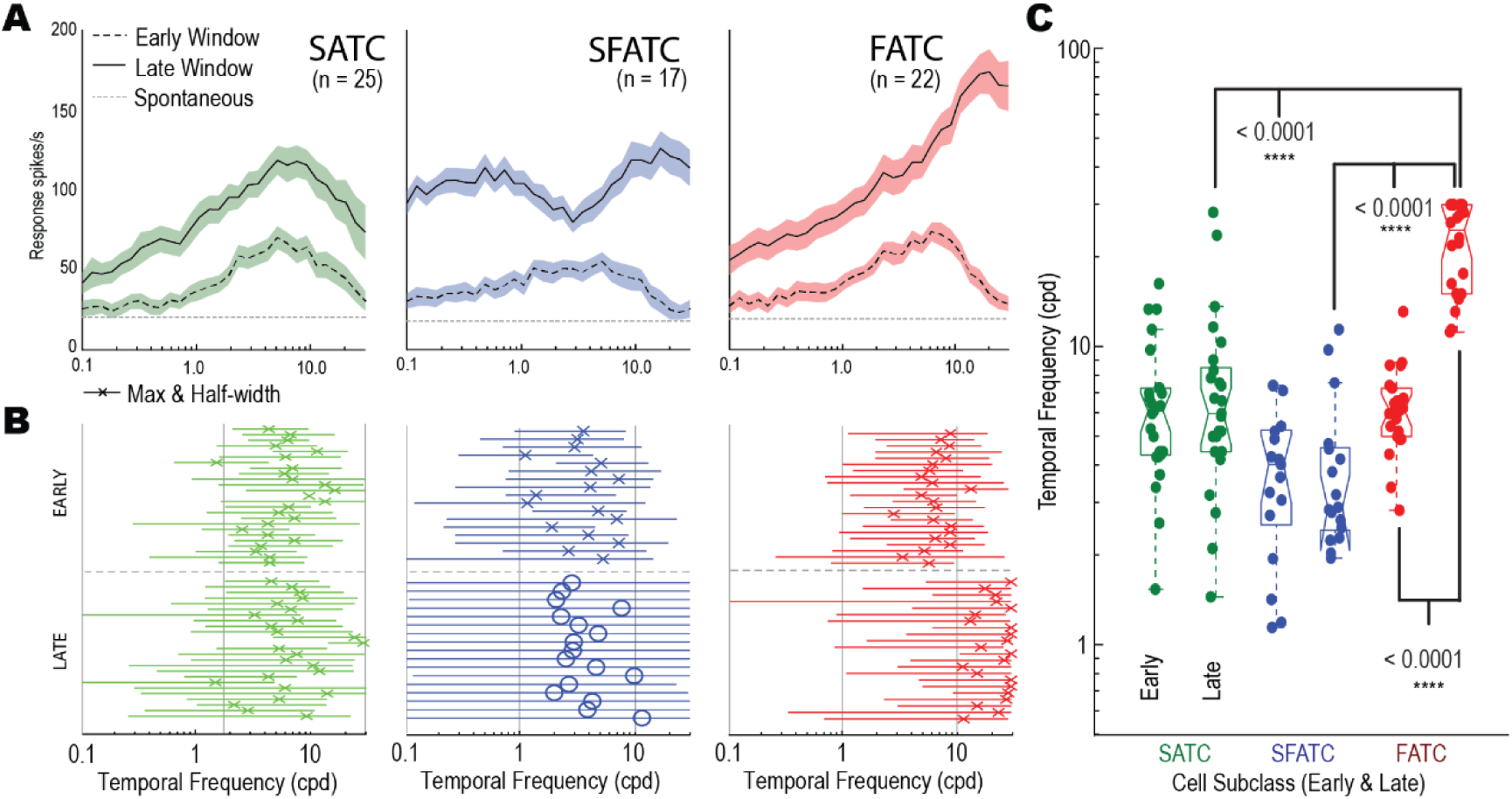
Temporal tuning of different LTC subclasses. **A**, Average temporal tuning for different cell classes in two response windows (early, late) as defined in Figure 4. Each plot shows the mean response as a line with the shaded area denoting standard error. FATCs exhibit a clear shift towards higher temporal frequencies in their late window compared with SATCs & SFATCs. **B**, Individual neuron summary data showing the position of the peak in early/late windows (denoted by an X) with the full width at half maximum (width of responses greater than 50% of maximum response) indicated by horizontal lines. In this case SATC cells exhibit little change in their tuning properties, FATC cells exhibit a large shift towards higher temporal frequencies and SFATC cells exhibit a characteristic adaptation at their former preferred temporal frequency. In (B), the position of the ‘notch’ in the adapted state is denoted by O rather than an X for the optimum for the SFATCs. **C,** Boxplot distributions for changes in temporal frequency optima. This metric cannot be defined in the late window for the SFATCs due to the notch. FATC neurons show a large increase in temporal frequency optima compared to their early window and a higher temporal frequency optima in the late window than both SATC and SFATC cells (using the notch location as a stand in for peak).

### 5.8 Temporal Tuning

Figure 8 shows the temporal tuning data (spatial frequency = 0.1 cycles/°) averaged across all cells allocated to the SATC, SFATC and FATC subclasses, using a similar analysis as for the spatial frequency domain (Figure 7). This largely supports our qualitative observations based on individual cells. The SATC temporal frequency tuning response is broad, with an optimum in both the early and late window of approximately 6 Hz. Apart from some broadening of the late window response (Figure 8A) the shape of the tuning function is very similar in both analysis windows.

SFATCs exhibit a pronounced change in temporal frequency tuning over the two windows measured. Reflecting their high contrast sensitivity, the temporal frequency tuning is already very broad in the early window, spanning more than 2 decades of temporal frequency (Figure 8A). This broadness is even more evident in the late window, with a prominent notch at intermediate temporal frequencies. This combines with additional recruitment of stronger responses to less optimal stimuli by higher contrasts to the full range of temporal frequency tested (0.1 to 30 Hz), such that the overall tuning is remarkably flat. Indeed the response never falls to below 50% maximum in this range in many cells, confounding our attempt to identify a distinct temporal optimum in the late window for these cells. This made it impossible to quantify either a change in the temporal optimum in the late window or a useful measure of response half-width (Figure 8B). Interestingly, if we identify the location of the notch in individual cells (indicated by ‘O’ symbols in Figure 8B) this coincides closely with the peak in the early window, suggesting that it arises from adaptation recruited more strongly at what was the initially optimal stimuli.

In SATC and FATC cells by contrast, a distinct maximum is seen in the response analysed in both early and late windows. In SATCs, the late window optimum is not significantly different than in the early window. In FATCs however we see a significant (at least a 4-fold) increase in the temporal frequency optima (Figure 8C, n=22, Kruskal-Wallis with multiple comparisons test). This new temporal frequency optimum was also significantly different to the temporal frequency optimum of the other two cell subtypes (Figure 8C, SATC n=25, SFATC n=17, FATC n=22, Kruskal-Wallis with multiple comparisons test). Note that due to the frame rate of our stimulus display, we limited stimuli to below 30 Hz in order to ensure that the phase shift for gratings on successive video frames never exceeded 90 degrees. In individual FATCs, the response is still rising at this upper limit, so our data set most likely underestimates the true magnitude of this shift in tuning. It seems likely that had we used a display with infinite frame rate, the roll off in response above 30 Hz would be primarily determined by the limits of early visual processing.

### 5.9 Velocity coding by LTCs for natural scenes

Our analysis of response tuning using narrow-band sinusoidal gratings suggests that all 3 LTC classes use fundamentally similar spatial and temporal filters in their underlying motion detectors, evidenced by their similar spatial and temporal tuning in the early windows. In other insects, such optima for sinusoidal patterns provide robust predictions for the velocity range over which the same neurons respond to broad-band images, including natural scenes (Dror et al., 2000, Barnett et al., 2010). However, as the contrast ramps progress, the large differences in adaptation to the stronger motion stimuli among different LTC classes may have a substantial influence on responses to moving natural patterns.

To test this we estimated velocity tuning using prolonged exposure to motion for a suite of either six or an extended set of sixteen natural image panoramas, depending on the experiment recording duration (Figure 9A). The stimulus comprised a sequence of brief periods of test motion across a wide range of velocities, interleaved with a constant adapting stimulus, but always moved continuously in the preferred direction for the neuron (Figure 9B). The adapting periods are longer than the brief test speeds (500ms versus 200ms) to ensure that the adaptation state is kept reasonably constant at the start of each test period. The test velocities cycle through an ascending and descending order to evaluate any hysteresis that may reflect differential adaptation to the test pulses themselves.

**Figure 9:**
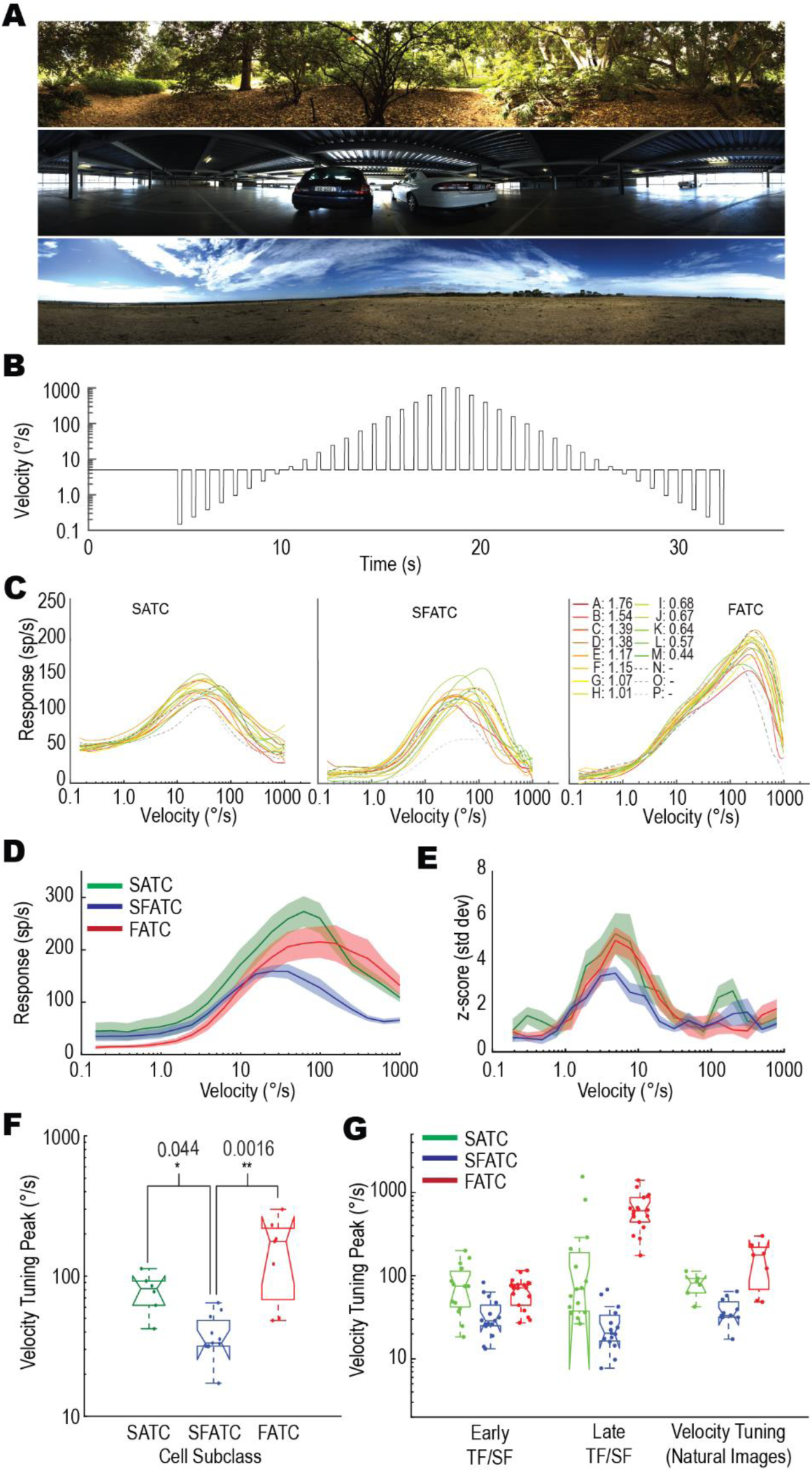
Velocity tuning of LTC responses to natural scenes. **A,** 3 examples from the set of 16 panoramic images used as stimuli. **B,** The velocity for each panoramic image is modulated over time, translated horizontally on the screen oriented in the preferred direction of the neuron. Following an initial 4 second period, brief test periods (200ms) of varying velocities were interleaved with longer (500ms) periods of the adapting speed (5°/s). **C,** Individual examples from three different cell subclasses, with each line a velocity tuning profile for a different natural image averaged across different starting phases and forward/backward sequences. Images A-P are color coded in order of their image contrast (C_EMD_). These LTCs exhibit consistency between images despite large changes in image contrast. In particular, note the consistency across images over decades of velocity for the example FATC. **D,** Responses of the three LTC subclasses to the test portion of the modulated velocity shown in (B) 50ms after the velocity change (100ms window, averaged across images/phases of forward and backward sequence). Each cell type exhibits a different velocity tuning profile with SATCs peaking at an intermediate velocity (62.8°/s), SFATCs peaking at lower velocities (24.5°/s) while FATCs peak at higher velocities (98.6°/s). **E,** A plot showing the z-scores (see text) at each velocity for each cell subtype (a measurement of information content. SFATC cells show the lowest information content**F**, Box plot showing the distributions of velocity optima for different LTC subtypes. P values indicate significantly lower optima in the SFATCs than the other two types (Mann-Whitney U). **G**, A box plot showing three estimates of velocity tuning for each of the three cell subtypes. Both early and late estimates use the tuning peaks calculated in Figure 7 and Figure 8 via the formula: Velocity = Temporal Frequency / Spatial Frequency. The final instead uses the late windows. The third shows the velocity tuning derived from natural images (Figure 9F). Velocity tuning from natural images falls between that calculated from the early and late windowed spatial and temporal tuning curves

For each neuron, we calculated the mean response across all image phases and produced a velocity-tuning curve for each background image. An exemplar of each cell subtype with 16 different images is shown in Figure 9C. As can be seen from these individual neurons, there is close agreement in response across images indicating that like LPTCs in Diptera (Straw et al., 2008), LTCs in dragonflies exhibit remarkable velocity constancy.

For each neuron, we then averaged the responses across images and grouped subtypes. We then plotted the mean and standard error for each subtype (Figure 9D, SATC n=6, SFATC n=11, FATC n=7). The motion adaptation we earlier described (Figure 8) manifests itself as differences in the optimal velocity tuning for each of the LTC subclasses. While all three subclasses of LTC begin responding to the moving background at ∼1°/s, their behaviour at fast velocities is markedly different. SFATC neurons exhibit the lowest velocity peak, occurring at °/s. SATC neurons are next, at 62.8 °/s. FATC neurons show the broadest tuning range peaking at 98.6 °/s. The ordering is in line with our previous results of temporal frequency analysis (Figure 8) indicating that the motion adaptation seen in the grating experiments translates into different velocity-tuning optima for natural images.

We also analysed the velocity constancy (lack of variation between images) seen in Figure 9C to see if there was a difference between the three subclasses. To assess the velocity constancy we used a metric, called a z-score. The z-score captures the response variation between different velocities (i.e. the useful information) compared to the response variation at a single velocity due to the change in background image. Z-score is defined here as:

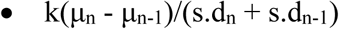

Where μ_i_ is the mean response (across images) at the i^th^ velocity and s.d_i_ is the standard deviation at the i^th^ velocity. The change in mean response between successive velocities (i.e. the information) is divided by the variation between images for a single velocity pair. This value is then normalized by a factor (k) to account for the number of velocity samples per decade of velocity (higher resolution samples show smaller mean differences between subsequent velocity measures). This final measure (z-score) gives an indication of which velocities ranges LTCs convey the most information (i.e. their dynamic range).

Figure 9E shows that FATC and SATC neurons exhibit a much higher z-score at their peaks than SFATC cells indicating that they convey more information about velocity (potentially improving the precision of the velocity estimate for example). Though subtle, the z-scores also seem to reiterate the slight difference in peaks between FATC & SATC neurons (with FATC being faster). Figure 9e show three individual examples of 16-image trials for a single neuron. Here we see the largest difference observed thus far. While the SFATC and SATC are qualitatively similar, the FATC neuron exhibits a significantly higher optimum velocity, around 150°/s, and a remarkably asymmetric velocity tuning function that rises almost monotonically over more than 2 decades of image velocity.

Figure 9F shows a boxplot of the peaks of the individual neurons separated by type. In agreement with Figure 9D both SATC and FATC cells have higher peak velocity than SFATC cells (SATC n=6, SFATC n=11, FATC n=7, Kruskal-Wallis with multiple comparisons). We did not find a statistically significant difference between peak of the the SATC and FATC neurons, though this is may be a consequence of insufficient power. We note the challenging nature of conducting intracellular recordings, particularly with respect to holding a neuron long enough to perform the velocity experiment across multiple images.

How do these velocity-tuning peaks relate to the temporal and spatial tuning of the neurons? To test this we calculated the predicted velocity tuning by dividing the temporal tuning peak by the spatial tuning peak (TF/SF) for both the early and late windows from Figure 7 and Figure 8 (Figure 9G). These were highly informative calculations. As can be seen from the boxplots, the velocity tuning peaks are in line with directions expected from temporal and spatial tuning peaks. The real velocity tuning data for FATC and SFATC neurons lies between the estimates derived from the early and late peaks. The SATC neurons appear fairly constant across all three methods of calculation.

**Table 1:** Tables showing the estimated velocity tuning peaks (median) of the three LTC subtypes. The estimates are as follows: 1, an estimate based on the early-window spatial and temporal tuning peaks, 2, an estimate based on the late-window spatial and temporal tuning peaks, 3, the peak calculated from our experiments in response to natural images of varying velocities (Figure 9)

|  | TF/SF Early (°/s) | TF/SF Late (°/s) | Natural Images (°/s) |
| --- | --- | --- | --- |
| SATC | 74.5 | 68.8 | 81.3 |
| SFATC | 28.7 | 20.3 | 33.3 |
| FATC | 71.2 | 602.5 | 176.6 |

## 6 Discussion

### 6.1 Effect of adaptation on velocity tuning

Overall, our results show some similarities between the dragonfly LTCs and their Dipteran counterparts, but also a number of intriguing differences. In Dipteran LPTCs such as the H1 neuron of the blowfly for example, previous work suggests that exposure to motion at high temporal frequency (20 Hz) leads to a rapid decay in response compared with the initial level, while at 1 Hz, responses are sustained for many seconds (Jung et al., 2011). We observe precisely the opposite effect in the FATC cells of the dragonfly, with the strongest recruitment of responses by the end of the stimulus at the highest temporal frequency tested. Does the apparent shift of maximal sensitivity to very high temporal frequencies and to very low spatial frequencies that we observed towards the end of the contrast ramps for these neurons reflect an adaptation mechanism that shifts maximal sensitivity from very low speed to very high speed? Such a mechanism has previously been proposed as an explanation for motion adaptation in different species, including humans (Clifford et al 1997). Contrary to the predictions of this model, however, subsequent estimates of temporal tuning before and after adaptation to a *constant* motion stimulus (typically a fast moving pattern) showed that the time constant of the underlying delay filter responsible for temporal tuning is not altered, at least in Dipteran LPTCs. Rather, adaptation primarily reduces the response via a decrease in the contrast gain of the neuron (Harris et al 1999, 2000).

In FATCs the very weak responses at low temporal frequencies late in the ramp, but very strong responses at high frequencies certainly suggests an adaptation mechanism that is more strongly recruited by slowly changing patterns in these cells – i.e. some form of potent non-linear high pass filtering recruited primarily by higher contrasts. However because the adapter is not constant in each trial, we cannot directly infer from our data that the delay time constant of the motion detector itself has changed. Nevertheless, since the contrast of natural scenes is also very high and dragonflies experience prolonged exposure to motion across a large range of velocities during natural behaviour, it is interesting to consider how this complex adaptation interacts with the underlying spatiotemporal tuning to shape velocity tuning during natural image motion among the different neuron subclasses.

While the higher velocity optimum in the FATCs would at first seem to be correlated with a difference in adaptation that leads to a higher optimum in the temporal frequency tuning and corresponding lower optimum in spatial frequency in our late window ramp data, more careful evaluation of the theoretical relationship between responses to narrow band sinusoids and broad band natural scenes would suggest otherwise. If we first consider the spatial and temporal optima from the early window in ramp data around 0.1 cycles/° and 6 Hz respectively for FATCs, these would predict an optimum speed of ∼60°/s for such narrow band sinusoidal patterns (Figure 9G). Natural scenes, however, have their highest contrast at low spatial frequencies, with contrast then declining at higher frequencies according to the famous 1/f^n^ characteristic (Field 1987, Tolhurst 1992). This 1/f property leads to velocity optima for such scenes a little over 2x those predicted for sinusoids for the same motion detectors (Dror et al., 2000). In other words, the observed velocity optimum for the FATCs in response to natural images, at around 120°/s is actually a good match for the predictions based on the spatial and temporal tuning in the unadapted (early) windows. Indeed, both the temporal and spatial tuning optima for sinusoids and the optimum speed for natural scenes for these dragonfly neurons are similar to those observed in LPTCs of male hoverflies (Barnett et al 2010, Straw et al 2008).

Hence it is actually the very slow velocity optima of the SATC and SFATCs, around 1/3 that of the FATCs, that is surprising. These optima are inconsistent with the predictions based on their responses to low contrast sinusoids, despite the latter being similar across all 3 groups (Figure 8C, Figure 9C, Figure 9G). This suggests that motion adaptation or other non-linear processing during prolonged exposure to motion of natural scenes leads to suppression of responses to higher velocities, which otherwise ought to be a more potent stimulus for these two cell classes. For SFATCs at least, this conclusion is consistent with the appearance of the ‘notch’ that we observe in the temporal frequency tuning for initially optimal patterns. Full resolution of this issue will require extensive future work employing a rigorous test-adapt-test approach to examine how the different components of motion adaptation previously identified in dipteran LPTCs (Harris et al 2000) are differentially recruited by different adapting stimuli.

### 6.2 Alternative Explanations of Motion Adaptation

Previous studies have demonstrated that differences in early versus late responses to motion at different temporal frequencies can be induced by differences in the behavioural state of the animal (e.g. during tethered flight versus restrained states) (Chiappe et al., 2010; Maimon et al., 2010). Many features of this differential adaptation to prolonged stimuli can also be induced in restrained animal preparations by exogenous application of agonists for the neuromodulator octopamine, and lead to apparent shifts in the temporal tuning of Dipteran LPTCs (Jung et al., 2011; Arenz et al., 2017; Suver et al., 2012). Could differences in octopaminergic modulation of the different classes of dragonfly LTCs potentially explain some of the differences we observe in their response time courses? Although we did not attempt to address the role of octopamine directly, it seems unlikely to explain the unusual adaptation we observe in FATCs for several reasons. Firstly, it was not uncommon to record data for two or more subclasses of LTC in the same animal exhibiting very different motion adaptation. This argues strongly against global differences in octopaminergic activity as an explanation. Secondly, either increased locomotor activity or octopamine agonists appear primarily to downregulate LPTC response *reduction* at high temporal frequencies during prolonged motion exposure (Jung et al., 2011). Hence the higher gain seen at higher temporal frequency following application of octopamine agonists in dipteran LTPCs results primarily from less adaptation to motion during the response time course. By contrast in FATCs we already see sustained, vigorous responses at the highest temporal frequencies tested, despite our animals being fully restrained. Instead we see selective adaptation for lower temporal frequencies and high spatial frequencies: both are more consistent with some form of powerful redundancy reduction in the signals at the elementary motion detector inputs. Finally, we have individual recordings where the temporal tuning was repeatedly measured that lasted over four hours with no apparent change in response shape.

Strong temporal and spatial (centre surround) antagonism have both been observed in dragonfly lamina monopolar cells (Laughlin 1974). Although linear spatial filtering ought to equally affect the early response window of our ramp stimuli, it would hardly be surprising if potent antagonism – either spatial or temporal – were recruited non-linearly as contrast increases, e.g. via additional voltage gated or inactivating conductances in feed-forward synapses. Indeed, such a mechanism may be required for a system with high contrast sensitivity (as observed here) to regulate the gain of local motion detectors and limit saturation in the real world, where average contrasts are high. Hence the differences we see between LTC subclasses may potentially arise from differences in which classes of lamina cells (or their post-synaptic targets) lie on the inputs to underlying motion detectors, an observation further supported by the differences we see in their transient responses to flicker stimuli. Note that both strong antagonism and non-linear temporal high pass filtering are key components of models proposed to explain spatiotemporal tuning of local motion detecting elements for the small target motion detector (STMD) pathway involved in target tracking in the lobula in these same dragonflies (Wiederman et al 2008), so it is possible that the SFATC and SATC subtypes of STC take their primary inputs from the same local motion detectors as these feature-detecting neurons.

### 6.3 Velocity constancy for natural scenes

In all 3 LTC classes, we observed a high degree of consistency across responses to natural images, with the majority of the curves peaking at a similar optimal velocity and the gain in response to different velocities being similar despite very large differences in the global contrast among this set of test images (Barnett et al 2010, Brinkworth & O’Carroll 2009). Such ‘velocity constancy’ for highly variable natural scenes in dipteran LPTCs has previously been shown to derive from a number of dynamic non-linear processing stages in biological vision, commencing with fast temporal adaptation in the photoreceptors and second order neurons, but also with a strong contribution from dynamic gain control within the local motion detectors (Shoemaker et al 2005, Brinkworth & O’Carroll 2009). Whatever the underlying mechanisms responsible for this impressive velocity constancy, some of the neurons recorded (particularly FATCs) come closer to being ideal velocity estimators than anything previously described at a single neuron level, in any animal, with a progressive, monotonic rise in response over more than a 100-fold range of velocities (Figure 9C). The consistency of these responses over that range is especially remarkable considering the enormous range of global contrasts observed in the set of images used in this experiment, as described in previous papers employing the same image set (Brinkworth & O’Carroll 2009)

### 6.4 Behavioural Implication

Hemicordulia dragonflies exhibit numerous distinct behaviours including hawking, patrolling and aerobatic conspecific engagements. Each of these tasks places different constraints on any system encoding optic-flow information and this would provide selective pressure on dragonflies to either adopt an extremely flexible motion-detection system or one which specialised for different tasks. Here we have shown that Hemicordulia have several different systems for encoding different kinds of ego-motion. The question is how the different LTC subclasses lend themselves towards the dragonfly’s behavioural tasks. In hawking behaviour, the detection and cancelling of subtle perturbations due to airflow requires a motion system capable of detecting slow velocities. SFATC cells exhibit a surprisingly robust response to grating patterns even at the very sluggish 0.1 °/s waveform. Such neurons would appear to be perfectly situated in a system designed to detect the slow-sustained shifts in optic flow that might occur during hawking behaviours.

Meanwhile, FATC cells exhibit extremely strong adaptation to slow-motion and are better suited to fast moving tasks such as patrolling or conspecific encounters. These neurons also show a preference for low spatial frequencies which is a more important background feature in fast-moving engagements. These neurons exhibit exquisite velocity constancy in natural image experiments despite great variations in the background, similar to those found in flies (Straw et al 2008).

Finally, SATC cells exhibit similar properties to LPTC found in flies, which have been linked to turning behaviours such as the optomotor response (Haikala et al., 2013). Unlike SFATCs, SATCs exhibit more classical tuning showing weak responses at both high and low temporal frequencies. In concert with FATCs, they may simply extend the velocity ranges over which Hemicordulia can operate (i.e. a slow and fast motion detector). As tuned neurons they are well capable of describing the magnitude of turning (ambiguities being eliminated using the faster FATC neurons) and thus could be used for relatively slower manoeuvres.

